# Bayesian optimization of separation gradients to maximize the performance of untargeted LC-MS

**DOI:** 10.1101/2023.09.08.556930

**Authors:** Huaxu Yu, Puja Biswas, Elizabeth Rideout, Yankai Cao, Tao Huan

**Affiliations:** Department of Chemistry, Faculty of Science, The University of British Columbia, Vancouver Campus, 2036 Main Mall, Vancouver, BC, Canada V6T 1Z1; Department of Cellular and Physiological Sciences, Life Sciences Institute, The University of British Columbia, Vancouver Campus, 2350 Health Sciences Mall, Vancouver, BC Canada V6T 1Z3; Department of Chemical and Biological Engineering, The University of British Columbia, Vancouver Campus, 2360 East Mall, Vancouver, BC Canada V6T 1Z3

## Abstract

Liquid chromatography (LC) with gradient elution is a routine practice for separating complex chemical mixtures in mass spectrometry (MS)-based untargeted analysis. Despite its prevalence, systematic optimization of LC gradients has remained challenging. Here we develop a Bayesian optimization method, BAGO, for autonomous and efficient LC gradient optimization. BAGO is an active learning strategy that discovers the optimal gradient using limited experimental data. From over 100,000 plausible gradients, BAGO locates the optimal LC gradient within ten sample analyses. We validated BAGO on six biological studies of different sample matrices and LC columns, showing that BAGO can significantly improve quantitative performance, tandem MS spectral coverage, and spectral purity. For instance, the optimized gradient increases the count of annotated compounds meeting quantification criteria by up to 48.5%. Furthermore, applying BAGO in a *Drosophila* metabolomics study, an additional 57 metabolites and 126 lipids were annotated. The BAGO algorithms were implemented into user-friendly software for everyday laboratory practice and a Python package for its flexible extension.

---

Liquid chromatography-mass spectrometry (LC-MS) is a sensitive and high throughput analytical solution that has been widely used for untargeted chemical analysis in proteomics^1, 2^, metabolomics^3, 4^, lipidomics^5^, and exposomics^6^, demonstrating great performance in explaining living processes from the chemistry level^7, 8^. In this technique, LC plays a vital role by separating compounds in the sample mixture, which significantly benefits the MS measurements by reducing ion suppression^9^ and co-fragmentation of isobaric species.^10^

Given the diverse chemical composition of samples, LC is usually operated with gradient elution. This technique facilitates the rapid separation of compounds with varying polarities, resulting in enhanced chromatographic peak resolution compared to isocratic elution.^11, 12^ To obtain high-quality MS data, LC gradient needs to be optimized to minimize compound coelution. Traditional design-of-experiment (DOE) starts with a user-defined satisfactory gradient and explores whether a similar gradient might be better.^13, 14^ Due to the substantial search space of potential gradients (typically exceeding 10^5^, **Supplementary Note 1**), conventional DOE lacks thorough exploration and its effectiveness heavily relies on the user’s initial gradient input. As such, DOE strategy is not widely used for LC gradient optimization. In fact, most gradient configurations are often under-optimized in LC-MS-based untargeted chemical analysis. Researchers tend to use a linear gradient or roughly adjust the gradient based on experience without a comprehensive performance evaluation. We advocate for the development of an optimization strategy that holistically considers all viable gradients while upholding efficiency, to systematically enhance LC separation power.

Bayesian optimization is a promising machine learning strategy for optimizing complex, black-box functions that are expensive and time-consuming to evaluate.^15^ It has found widespread use in hyperparameter optimization in machine learning, where evaluating a single set of hyperparameters requires significant computational resources for model retraining.^16, 17^ The advantage of Bayesian optimization lies in its ability to strike a balance between exploration and exploitation, focusing on areas with high expected outcomes while simultaneously probing regions with high uncertainty. This approach helps find the global optimum while minimizing the number of evaluations required for expensive experiments. In recent years, Bayesian optimization has found compelling applications in the field of chemistry, showcasing its promising performance in chemical synthesis, material design, among others.^18–23^

Here, we present BAGO, a dedicated Bayesian optimization framework and open-source software for LC gradient optimization. BAGO evaluates the retention of all detected features in an unbiased manner regardless of ion abundance and identity, providing a robust index representing global compound separation. Multiple optimizations of general Bayesian optimization framework were applied to ensure the high efficiency of BAGO on a diverse range of gradient optimization problems. As a fully automated approach, we believe it can be seamlessly integrated into routine analytical workflows requiring no coding experience from users. To ensure versatility and extensibility, an application programming interface (API) was developed as a Python package ’*bago*’.

## Results

### Development of BAGO

Bayesian optimization finds the optimal LC gradient through a sequential strategy (**Fig. 1a**). The process begins with the initial gradient to be optimized. By analyzing the LC-MS data obtained from the initial gradient, a new and promising gradient is predicted for validation through next experiment. If the proposed gradient yields unsatisfactory compound separation, the collected LC-MS data will be combined with previous data to recalibrate the subsequent gradient candidate. This sequential refinement strategy iterates until the paramount gradient is ascertained (**Fig. 1b**).

**Fig 1.**
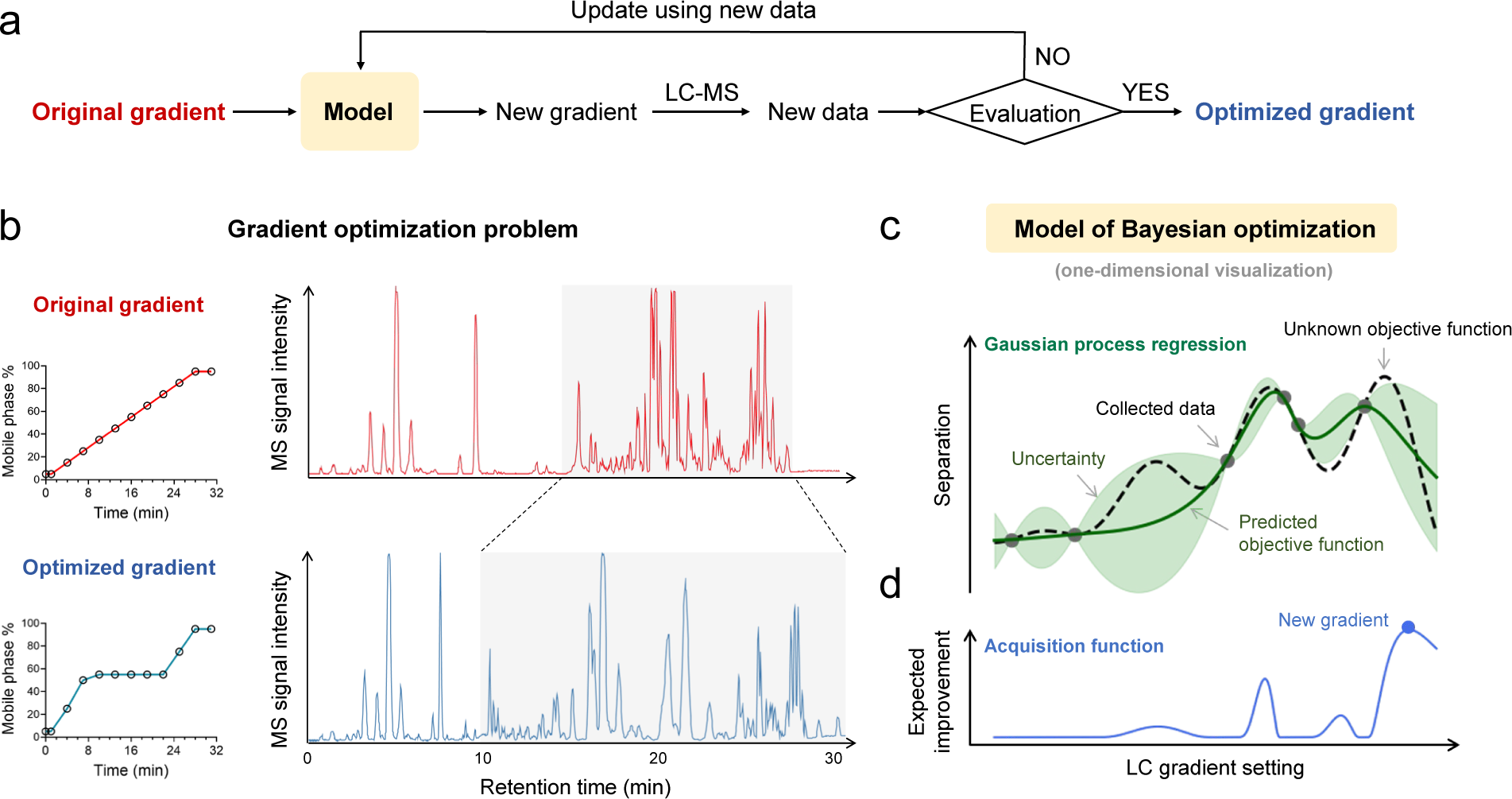
**a**, Flowchart shows general architecture of Bayesian optimization of LC gradients. **b**, Improved compound separation after optimizing LC gradient is visualized by base peak chromatograms. Data were collected from a human serum lipidomics sample separated on a reverse phase column in 31 minutes. The black circles represent the percentages of mobile phase B. **c**, One-dimensional visualization of the Bayesian optimization of an LC gradient. A Gaussian process regression (GPR) model predicts the unknown objective function with uncertainty. **d**, An acquisition function generated from GPR determines the next LC gradient to run.

The key step in Bayesian optimization is to predict the most promising gradient from observed data, which is achieved by “model” as shown in **Fig 1a**. In this step, a surrogate model is first constructed to approximate the unknown functional relationship between compound separation and LC gradient (i.e., unknown objective function). Gaussian process regression (GPR)^24^, a powerful surrogate model, is typically utilized in Bayesian optimization. Importantly, GPR predicts the unknown objective function with quantified uncertainty (**Fig. 1c**). A region of high uncertainty means it lacks observed data and has limited knowledge. To efficiently find the global maximum of the unknown objective function, a typical dilemma is to decide whether to explore the regions with high uncertainty (i.e., exploration) or to exploit the regions around the best observation (i.e., exploitation). Exploration maximizes the knowledge that can be gained in the next experiment regarding the unknown objective function, but it may also result in unnecessary effort spent on querying low-yield regions. On the other hand, exploitation typically ensures a promising outcome, but it risks getting trapped at a local maximum.

To balance exploration and exploitation, an acquisition function is created based on the result of GPR. **Fig. 1d** shows a common acquisition function termed expected improvement (EI). It comprehensively considers the GPR-predicted mean and uncertainty, balancing exploration and exploitation. The gradient with the highest value in the acquisition function will be tested in the next LC-MS experiment. As more data are collected and fed into the model, knowledge regarding the objective function accumulates, increasing the chance of finding the optimal gradient.

To apply Bayesian optimization on LC gradient, the first critical step is encoding, which transforms gradient configurations and compound separation performance into numerical values for downstream computation. In this study, a gradient configuration is represented as a *p*- dimensional vector, with each element denoting the mobile phase percentage at a specific time point (**Methods**). To assess separation performance, we proposed the global separation index (GSI), which is a singular value that evaluates the global compound separation (**Fig. 2a**). To compute GSI, all MS signals are first inspected, and then the peak apexes from unique compounds are selected as top signals (**Methods**). Next, a sequence of retention time intervals between adjacent top signals is squared and summed, defined as SQRTI (<u>s</u>um of the s<u>q</u>uared retention time intervals). SQRTI is a bounded value (**Supplementary Note 2**) that reaches the maximum with no separation and the minimum with perfect separation (i.e., all compounds are equally spaced). GSI is then derived by normalizing SQRTI to a fixed range from 0 to 1, where a higher GSI means better separation (**Fig. 2b**). Importantly, the scale of GSI is independent of the number of selected MS signals and total elution time, making GSI a universal metric for global separation performance.

**Fig. 2.**
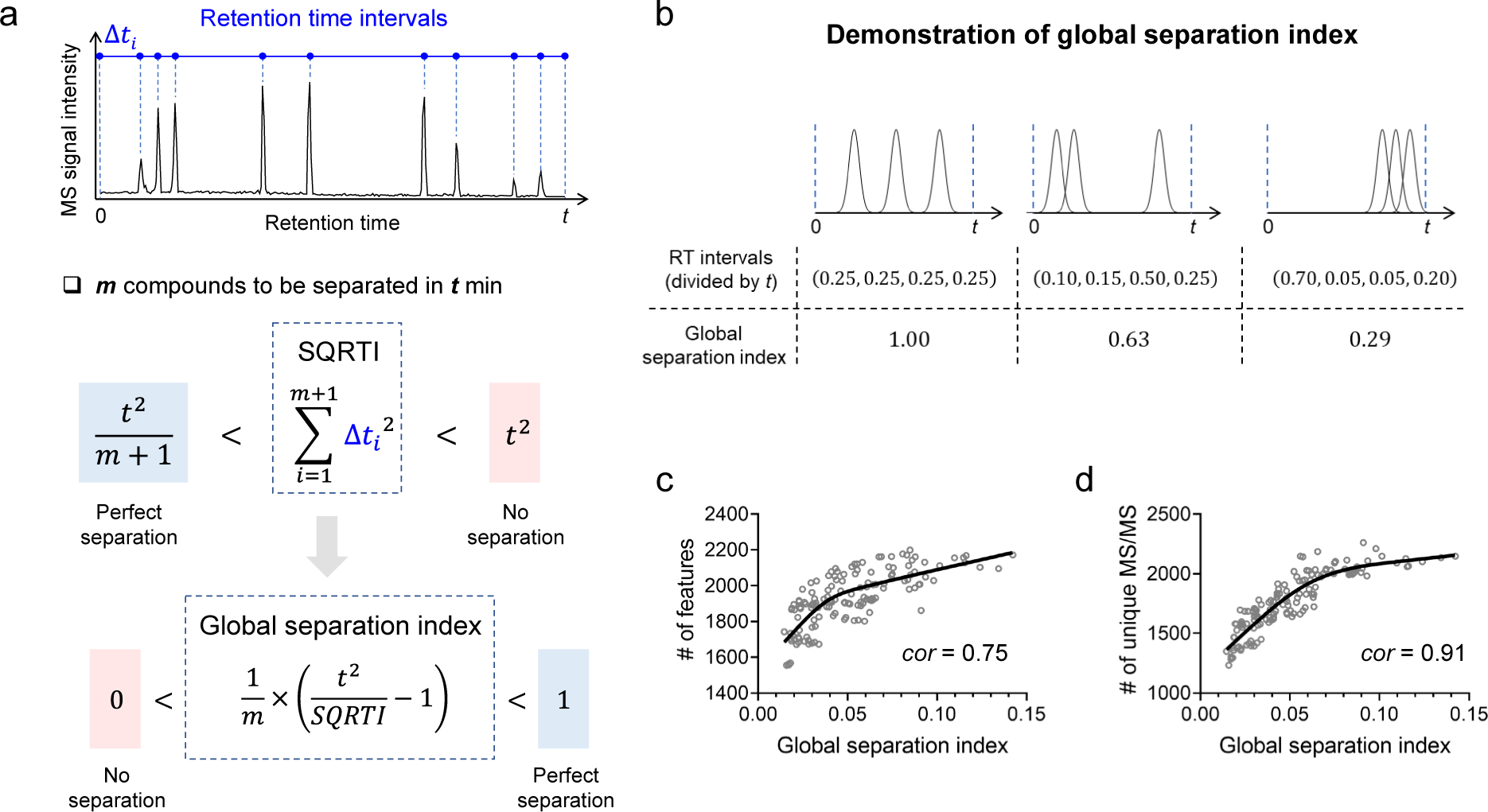
**a**, Encoding of omics-scale compound separation using a global separation index (GSI). SQRTI: sum of the squared retention time intervals. Perfect separation is defined as all compounds are equally spaced during the acquisition window; no separation is defined as all compounds eluted together at the beginning of gradient. **b**, Calculated GSIs of three visually different degrees of compound separation. **c**, **d**, GSI is highly correlated with the number of metabolic features (**c**) and number of unique MS/MS spectra (**d**). Grey circles represent 142 individual LC-MS/MS experiments with different gradient settings. Black lines were computed by fitting spline curves to show the general trend of data. *cor*: Spearman correlation.

We verified GSI as a reliable metric of LC separation performance by analyzing a human urine sample using 142 unique plausible gradient configurations (**Supplementary Note 3**). Among all tested configurations, GSI ranged from 0.0146 to 0.142, indicating that modifying gradient configurations can significantly change the global separation performance. In addition, high Spearman correlations of 0.75 and 0.91 were noted between the number of detected metabolic features (**Fig. 2c**) and unique MS/MS spectra (**Fig. 2d**), respectively, when compared against GSI. Optimizing the LC gradient based on GSI is demonstrated as a promising strategy for improving chemical detection and annotation with MS.

Following encoding of LC-MS experiment, we further refined the Bayesian optimization algorithm to maximize its efficiency for optimizing LC gradient. The algorithm efficiency is determined by acquisition function. Various acquisition functions have been proposed to balance data exploration and exploitation in different ways (**Supplementary Note 4**). Benchmarked on the total optimization steps, EI outperforms four popular acquisition functions to show the highest efficiency (**Extended Fig 1**. and **Supplementary Note 5**). Therefore, EI is used by default in BAGO, while other acquisition functions were also included in the *bago* Python package for implementation.

With the prepared encodings and algorithms, we present a comprehensive Bayesian optimization framework consisting of three stages: search space generation, initialization, and gradient optimization (**Fig. 3**). In the first stage, a search space is defined as a collection of plausible gradient configurations for an LC-MS experiment. These configurations adhere to two main primary constraints: (1) the percentage of the strong mobile phase should monotonically increase, following gradient design principles, and (2) the eluting power of a gradient, estimated by the total strong mobile phase used during the run, should remain within a reasonable range. By applying these constraints, gradients with poor separation or carryover issues are avoided. In the second stage, a Bayesian optimization model is initiated with two gradients specified by the user or selected by the algorithm. It is recommended to choose gradients with low correlation to emphasize initial data exploration. In the third stage, a GPR model is constructed based on the obtained data. Using the GPR model, an acquisition function is computed to identify a promising gradient (xnext) for the next evaluation, resulting in a new GSI value (ynext). This iterative process allows the GPR model to continually refine itself and eventually converge to the global optimal gradient. In practice, the rounds of optimization depend on the budget of time and resources. Our results, derived from four different gradient optimization problems, suggest that conducting ten rounds of optimization is sufficient to identify a satisfactory gradient (**Fig. 4**, **Supplementary Notes 6-8**).

**Fig. 3.**
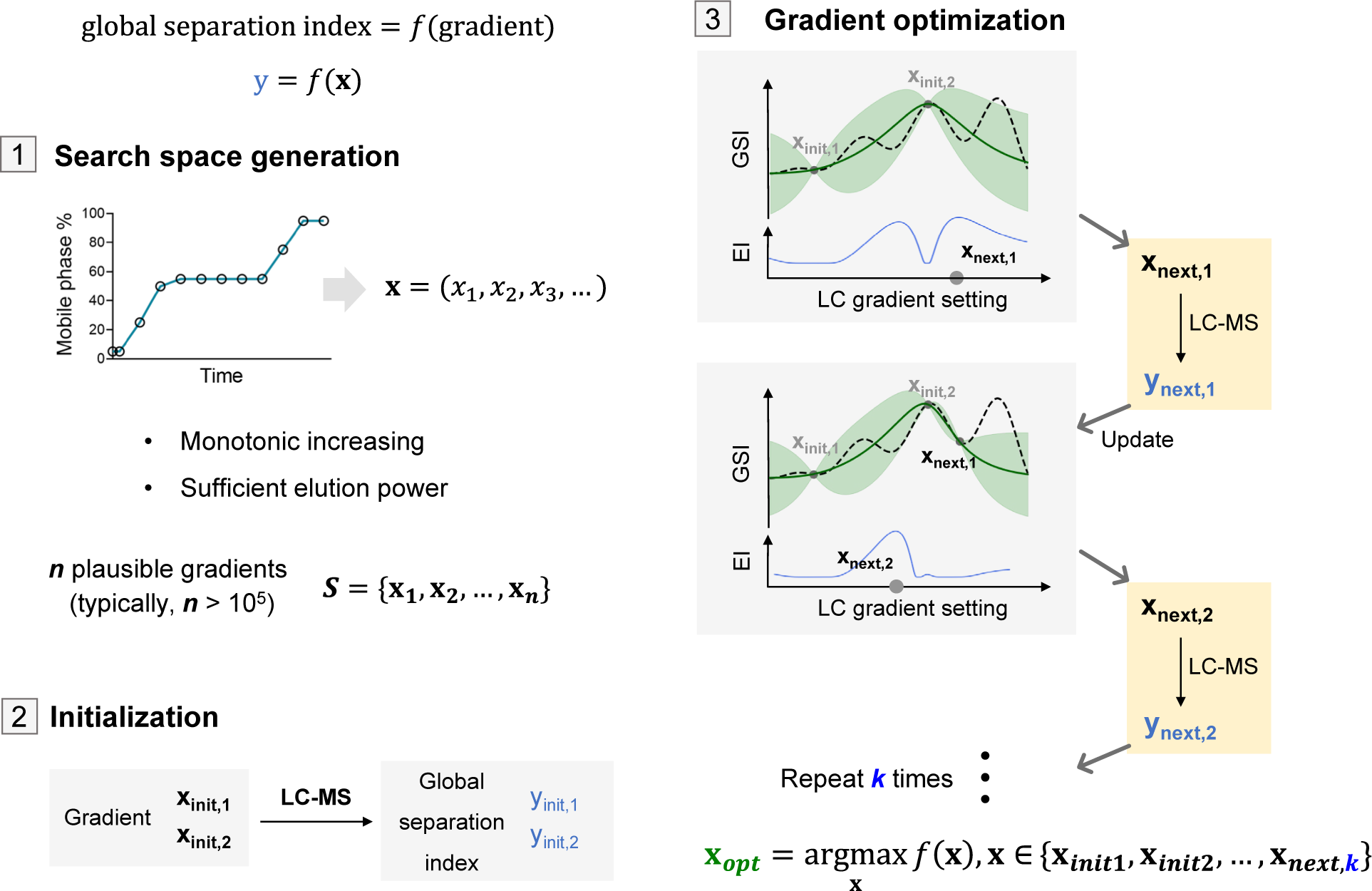
Schematic workflow of the Bayesian optimization of LC gradients that includes three stages: search space generation, initialization, and gradient optimization. The last stage, gradient optimization, is repeated by continuously taking new LC-MS data, updating the model, and providing a promising gradient for the next experiment. GSI, global separation index. EI, expected improvement.

**Fig. 4.**
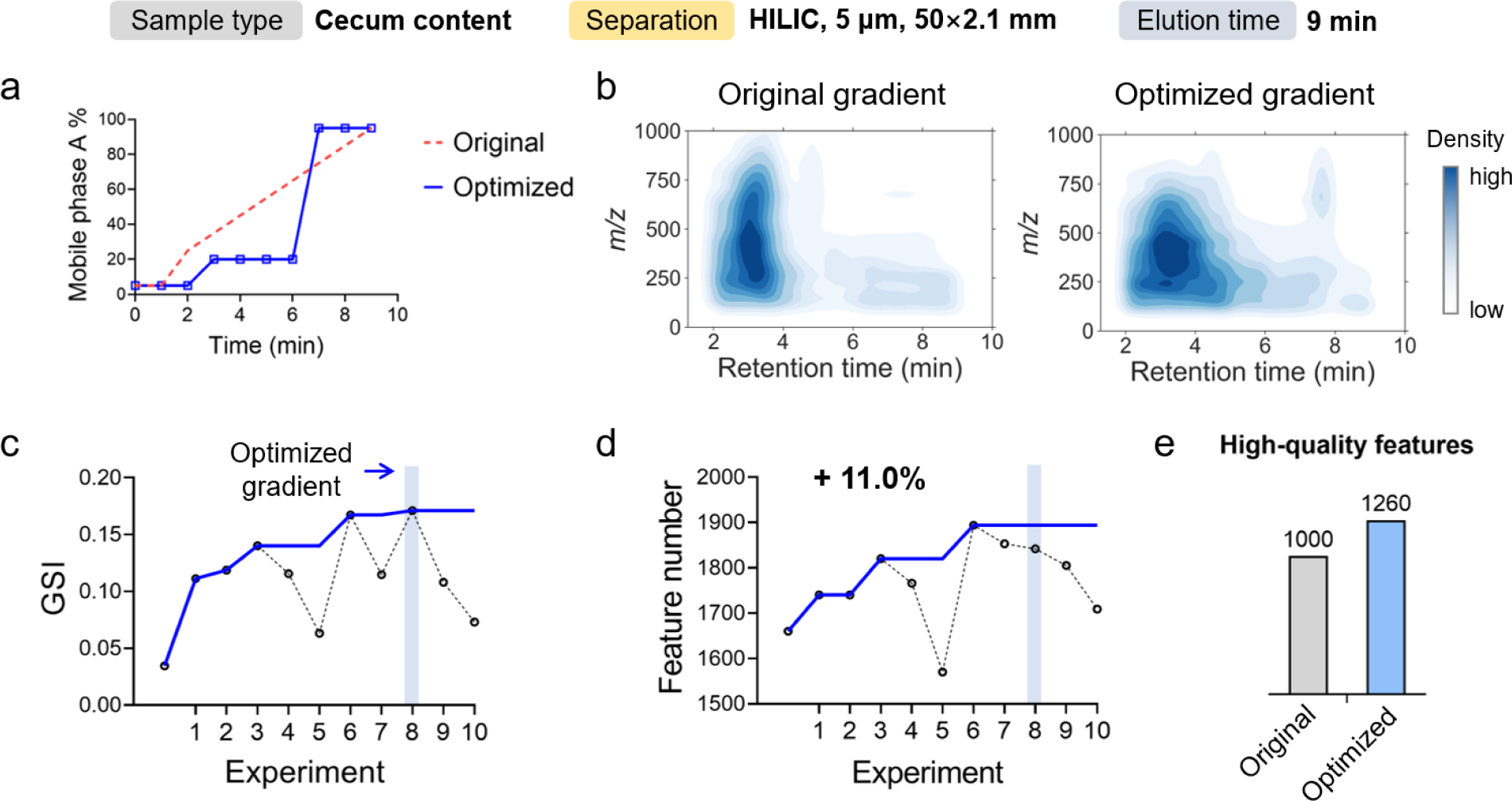
**a**, The optimized LC gradient vs. the original gradient. **b**, Improvement of compound separation visualized by a two-dimensional density plot of *m/z* to retention time. **c**, **d**, Improved global separation index (GSI) (**c**) and total number of metabolic features (**d**) during the 10- experiment optimization. The optimal gradient was found at the eighth experiment. Cumulative values were shown as solid curves. **e**, Improved number of high-quality features after gradient optimization. High-quality features represent the true metabolites with high quantitative accuracy and reproducibility, selected by applying multiple orthogonal criteria.

The entire data processing workflow was streamlined into user-friendly software with a graphical user interface (**Extended Fig. 2**). We also developed a Python API to support the proposed Bayesian optimization framework for customization, extension, and flexible implementation into other analytical pipelines. Besides, a YouTube video was created for its quick start guide (https://youtu.be/btNblKBXxk8)

### Performance

To validate the benefit of BAGO on improving LC-MS data quality, we performed an in-depth investigation using the mouse cecum metabolomics samples analyzed on a hydrophilic interaction chromatography (HILIC) column (**Fig. 4**). We began the optimization by defining a large search space that contains 261,484 plausible gradients. A ten-round optimization was carried out, starting from a simple linear gradient. We observed the best gradient in the eighth experiment. The optimized gradient reduced the climbing of the strong mobile phase from 0 to 6 minutes and dramatically increased it after. The strong mobile phase percentage was kept at 95% after 7 minutes until the end at 9 minutes to ensure sufficient elution (**Fig. 4a**). The density distribution of metabolic features on a two-dimensional graph (*m/z* versus retention time, **Fig. 4b**) visualizes the improvements in compound separation. In the original gradient, the high density of the region from 2 to 4 minutes indicates the gradient was increasing too fast, which was corrected in the optimized gradient. The optimized gradient improves the GSI from 0.0345 to 0.171 (**Fig. 4c**), leading to 11.0% more metabolic features (from 1660 to 1894, **Fig. 4d**). Our workflow increases the high-quality metabolic features by 26.0% (from 1000 to 1260) with satisfactory quantitative performance (**Fig. 4e**); these features were selected by applying multiple criteria^25^ to remove background ions, check analytical accuracy and reproducibility (see **Methods**).

Besides quantification, our method also facilitates compound annotation by improving MS/MS spectral acquisition. Using the same mouse cecum metabolomics data, we showed that the BAGO workflow improved the number of unique MS/MS spectra by 23.1% (from 1148 to 1413, **Fig. 5b**), indicating more metabolites can be annotated. Besides more MS/MS spectra, separating originally coeluted compounds reduces the number of chimeric MS/MS spectra, which were from co-fragmentation of different ion species and decreases the annotation accuracy^10, 26^. One example is shown in **Fig. 5a**. In the original gradient, two ions (ion 1: *m/z* = 130.0543 and ion 2: *m/z* = 132.0811) are highly coeluted with an *m/z* difference of 2.0268 Da. Even though the MS data were collected in data-dependent acquisition mode, the MS/MS spectrum of ion 1 was contributed by both ions. When searched against the MS/MS library, this convoluted spectrum failed to support the correct annotation of phenylacetylglutamine, showing a dot product similarity of 0.560. On the other hand, ion 2 was treated as the isotope of ion 1, excluded from MS/MS spectrum acquisition, and cannot be annotated. In the optimized gradient, these two ions were well separated with a 0.4 min retention time difference. Thus, clean MS/MS spectra were collected for both ions, leading to correct identifications and spectral similarities of 0.995 and 0.950 for ion 1 and ion 2, respectively.

**Fig. 5.**
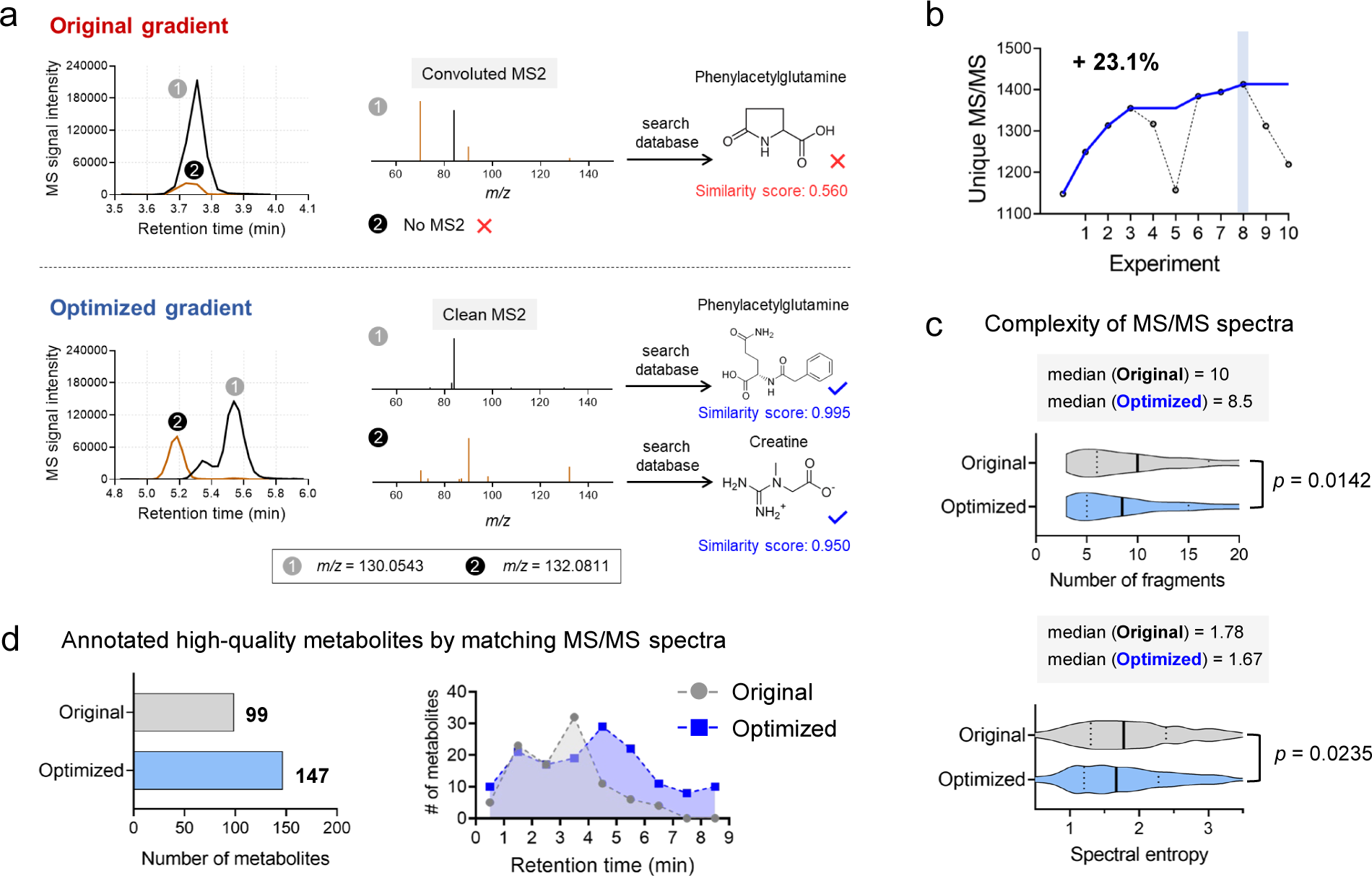
**a**, An example of gradient optimization facilitating compound identification by improving the coverage and quality of MS/MS spectra. **b**, Increase of unique MS/MS spectra during gradient optimization. **c**, Reduced spectra complexity by gradient optimization, characterized by number of fragments and spectral entropy. **d**, Increase of annotated metabolites after gradient optimization. Histogram shows the distribution of annotated metabolites over the 9 minutes of elution.

We further evaluated the improved spectral quality at the omics scale. The cleaner MS/MS spectra were evidenced by reduced spectral complexity. As shown in **Fig. 5c**, the optimized gradient showed significantly fewer fragments (median decreased from 1.78 to 1.67, *p* = 0.0235). The optimized gradient also scored a significantly lower spectral entropy, a value to index spectral complexity^27^, compared to the original gradient (median decreased from 1.78 to 1.67, *p* = 0.0235).

Of the high-quality metabolic features that fulfill the quantification criteria, the optimized gradient enables the annotation of 48.5% more metabolites compared to the original gradient (**Fig. 5d**). In a comparison of annotated metabolites distributed over retention time, we observed a similar distribution pattern before 3 minutes and a clear increase of annotations after 4 minutes.

The performance of BAGO was further validated on three more gradient optimization problems, including human urine metabolomics, serum metabolomics, and serum lipidomics (**Supplementary** Fig. 2-4). The robust improvement of compound separation by BAGO is characterized by GSI and visualized by two-dimensional graphs (*m/z* versus retention time). We observed that the number of detected features did not significantly change over 10%, yet a total of 13.0%, 10.6%, and 16.3% more unique MS/MS spectra were acquired in these three studies, respectively. Altogether, our results highlight that the proposed Bayesian gradient optimization strategy effectively enhances separation, facilitating untargeted chemical detection, quantification, and annotation.

### Biological Applications

We next demonstrated the BAGO workflow on *Drosophila* male and female abdominal carcasses using a parallel metabolomics and lipidomics workflow (**Supplementary Note 9**). While prior studies have begun to determine how genetic variation and diet influence *Drosophila* metabolites or lipids using single- or mixed-sex animal groups^28–32^. However, a comprehensive and sex-based analysis of lipids and metabolites has not been completed. Defining sex differences in metabolites and lipids can offer vital insights into the sex-biased risk of developing metabolic dysregulation and disease across multiple animals^33–36^. In this study, we designed an untargeted metabolomics analysis using HILIC separation with an 8-min gradient and a lipidomics analysis using RP separation with a 25-min gradient (**Fig. 6a**). Ten rounds of Bayesian gradient optimization were applied for each mode. As shown in **Fig. 6b**, we observed that the optimization increased GSI from 0.0267 to 0.0990 for metabolomics analysis and 0.105 to 0.197 for lipidomics analysis. Benefiting from the increased compound separation, we obtained 50.1% more (441 to 662) high-quality metabolic features and annotated 34.5% more (57 out of 165) metabolites. For the lipidomics analysis, we acquired 36.9% more (1268 to 1736) high-quality lipidic features and annotated 20.8% more (126 out of 606) lipids.

**Fig. 6.**
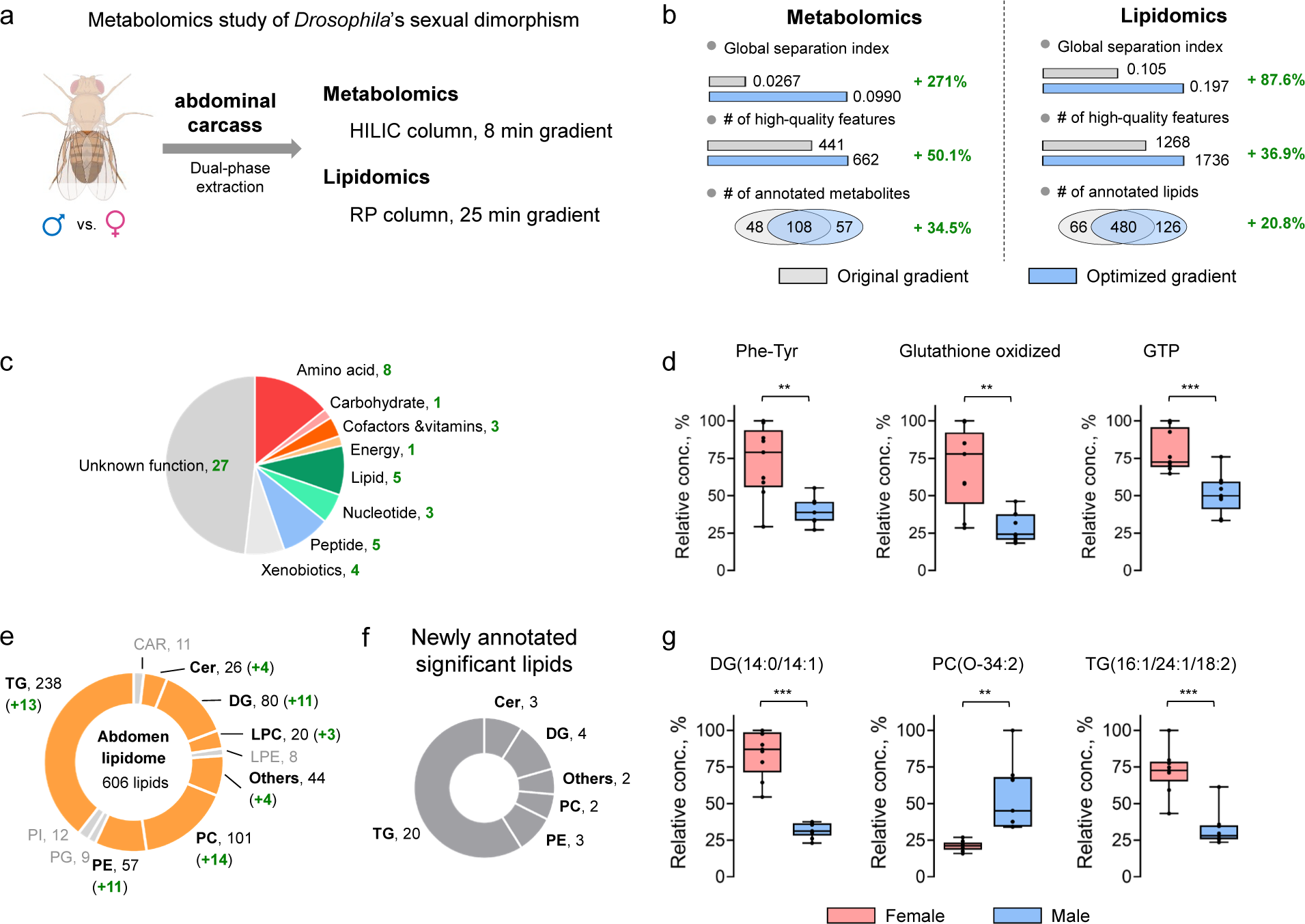
**a**, Experimental design of a parallel metabolomic and lipidomics study of *Drosophila*. **b**, Increase of global separation index, number of high-quality features, and number of annotated compounds after gradient optimization. **c**, Newly-annotated metabolites using optimized gradient. **d**, Box plots shows three newly confirmed significant metabolites between females (left) and males (right). GTP, guanosine triphosphate. **e**, Newly-annotated lipids classified into 11 classes. The numbers in brackets represent the increase from optimization. **f**, Newly-annotated significant lipids in six classes. **g**, Box plots shows three newly confirmed significant lipids between females (left) and males (right). DG, diglyceride; PC, phosphatidylcholine; TG, triglycerides. **: *t*-test *p* < 0.01, ***: *t*-test *p* < 0.001

With the optimized gradients, we further examined the metabolome and lipidome profiles for differences between males and females. For lipidomics results, we classified the detected lipids into ten main classes based on LIPID MAPS^37^ (**Fig. 6e**). Of these ten, eight classes show more annotations, such as phosphatidylcholine (PC, 14 more), phosphatidylethanolamine (PE, 11 more) and diglyceride (DG, 11 more). Comparing the males and females, we observed 34 newly annotated lipids with significant differences (*t*-test *p* < 0.05, **Fig. 6f**). Three of the newly annotated lipids, DG (14:0/14:1), PC (O-34:2), and TG (16:1/24:1/18:2), were highlighted through box plots in **Fig. 6g**. For the metabolomics study, we examined a total of 57 newly annotated metabolites and classified them into nine categories based on their main metabolism involvement from the KEGG pathway database (**Fig. 6c**). Eleven high-quality metabolites show a significant difference between males and females, including the Phe-Tyr dipeptide, glutathione oxidized, and guanosine triphosphate (**Fig. 6d**).

## Discussion

This work presents a Bayesian optimization framework that automates the optimization of LC gradients. We aim to transition the conventional experience-based LC experimental design to a data-driven approach, making the entire optimization workflow more automatic, reproducible, and feasible. Unlike conventional human decision-making strategies, our approach eliminates the need for manual interpretation of large, high-dimensional LC-MS data and does not rely on prior knowledge of analytes’ chemical structures. The proposed approach significantly improves the efficiency and robustness of global compound separation, leading to better GSIs ranging from 81.2% to 396% across six scenarios, each differing in biological sample type and LC column. Better chromatographic separation further benefits compound annotation with more and cleaner MS/MS spectra acquired. It also improves compound quantification by minimizing the MS signal interference among coeluted compounds. Its application to a *Drosophila* abdomen metabolomics study on both sexes demonstrated a noticeable increase in high-quality metabolic and lipidic features of 50.1% and 36.9%, respectively. This substantial increase leads to broader biological knowledge, acquired by using the BAGO workflow for gradient optimization. We implemented BAGO into a desktop application that requires no coding experience or chemistry knowledge for optimizing a gradient. We also provide a Python API for programming usage and to encourage contributions from the community for further development.

A fundamental challenge of gradient optimization in untargeted chemical analysis is the lack of a global compound separation metric. The previous work GOAT is a computational tool that optimizes LC gradients in proteomics. It aims to equally distribute the MS/MS spectra with the top 50% total intensities.^11^ However, the improved peptide separation was only visually supported by the base peak chromatogram and indirectly verified by the improved peptide and protein identification. Also, the gradient can be optimized based on retention time prediction given a set of known compounds. Hence, an *in silico* gradient optimization method was proposed for reverse phase separation in proteomics research.^38^ It relies on the prediction of compounds’ retention times using a specific gradient. Nevertheless, retention time prediction relies on prior knowledge of the molecular structures and is not suitable for untargeted analysis where a majority of chemicals are unknown^39^. Therefore, a metric considering all MS signals and independent of ion identity is highly desired.

Therefore, we proposed GSI as a robust metric of compound separation performance. GSI is calculated based on observed MS signals rather than only known chemicals, making it better suited for handling the numerous unidentified molecules in MS analysis (e.g., metabolomics). In addition, GSI is computed solely based on MS1 spectra; therefore it is independent from MS/MS acquisition and applicable to MS data acquired under full-scan, data-dependent, and data-independent acquisition modes^40^. Notably, the proposed GSI concept can be extended to other chemical analysis platforms coupled to LC such as ultraviolet–visible spectroscopy^41^ and electrochemical detection^42^, where the data structure of the chromatographic peaks is the same as MS.

Bayesian optimization is an active learning approach that searches for the most promising gradient by balancing data exploitation and exploration via an acquisition function. The success of a Bayesian optimization model thus relies on the performance of the acquisition function. In our strategy, expected improvement (EI) was selected as the default acquisition function with the smallest variance and worst-case loss and the fewest optimization steps (**Extended Fig. 1, Supplementary Note 5**). EI queries the chance of obtaining a higher outcome in the entire search space by considering the predicted mean and variance in the Gaussian process regression (GPR). As a robust acquisition function, EI was also used as the default in other Bayesian optimization frameworks such as EDBO for chemical synthesis^22^. In our study, EI outperforms the pure exploitation and exploration algorithms that only consider the predicted mean and variance, respectively (**Extended Fig. 1**). Notably, a hyperparameter *δ* in EI can be further tuned to emphasize exploration or exploitation. In our method, *δ* = 0.01 was set as default since it has been proven to deliver great optimization performance in a broad range of optimization scenarios^24^. We also confirmed that *δ* = 0.01 provides the highest optimization efficiency by testing it on a urine metabolomics data set.

The benefit of optimizing LC gradients is profound and can substantially improve the performance of LC-MS analysis beyond just resolving the closely eluting compounds. By reducing coelution, higher ionization efficiency is achieved, leading to more metabolic features that can be detected. This phenomenon has also been observed in proteomics; improving compound separation has been evidenced in identifying more proteins.^11, 43^ Moreover, the coelution of compounds with small *m/z* differences at the level of Daltons can be minimized with an optimized LC gradient. Less coelution benefits two aspects of the downstream metabolite annotation. First, the coverage of MS/MS acquisition was improved (**Fig. 5b**), leading to more unique MS/MS spectra available for compound annotation. Secondly, the chimeric MS/MS spectra are reduced (**Fig. 5c**), improving the spectral similarity of true identification when matching against an MS/MS spectral library. Even though bioinformatic strategies have been developed to deconvolute the chimeric spectra for improving identification accuracy^10, 44, 45^, we believe that reducing the acquisition of chimeric MS/MS spectra in the first place avoids the risk of false deconvolution and simplifies the entire process. With the enhanced quantification and identification, we achieved 48.5% more annotated metabolites that have high confidence in the mouse gut metabolomics study for further statistical analysis.

The proposed Bayesian workflow for gradient optimization works for a wide range of biological applications with different LC columns and sample types. Demonstrated on a comprehensive metabolomics and lipidomics study of *Drosophila* abdominal carcasses, BAGO shows it can boost the detection and quantification of both polar and nonpolar chemical compounds with different separation mechanisms. Therefore, it may serve as a useful tool for routine multi-modal untargeted analyses of small molecules. Overall, the development of this Bayesian optimization strategy enables highly efficient optimization of LC gradients for enhanced compound profiling in MS analysis. This approach has the potential to be seamlessly integrated into the control systems of LC-MS platforms, enabling fully automated gradient optimization without the need for human intervention^46^. By leveraging this proposed Bayesian optimization framework, we believed that rapid method development for omics-level biological and pharmaceutical research can be achieved, thereby expanding the scope of small molecule discovery and exploration.

## Methods

### Encoding LC gradient and separation performance

We consider the gradient optimization problem as finding the LC gradient setting (input) to achieve the best compound separation (output). LC gradient setting and global separation index are encoded. For each LC gradient, we used a linear vector to specify the mobile phase percentages at different time points. Here, the mobile phase with a monotonically increasing percentage (i.e., strong mobile phase) during an experiment was encoded, while the percentage of weak mobile phase can be inferred. Suppose that during the experiment, the encoded mobile phase ratio can be tuned at *n* time points. We defined an LC gradient setting **x** as

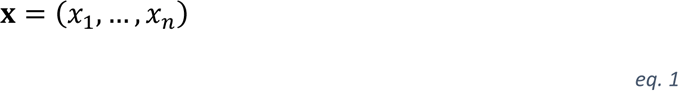

where *xi* represents the mobile phase ratio at the *i*th time point.

With the vector descriptor of LC gradient settings in hand, a search space consisting of all gradient settings to be tested was generated. To begin, a set of evenly spaced mobile phase percentages were created by defining the lowest and highest percentage and gradient step size. For instance, to find the best gradient of a strong mobile phase ranging from 30% to 70%, researchers may set the step size as 10% to obtain an array of five elements (30%, 40%, 50%, 60%, and 70%). Then, each *xi* in **x** is randomly selected from the array to form a gradient. Notably, two restrictions were applied to the search space generation. First, elements in each **x** are monotonically increasing to meet the empirical requirement of LC gradient design (**eq. 2**).

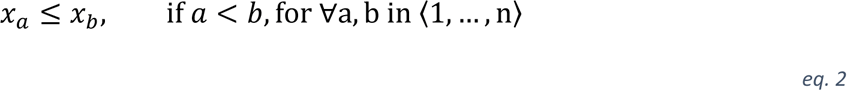

Second, we considered the overall portion of the strong mobile phase. It was restricted to a user-defined range to ensure the gradient is fast enough for all metabolites to elute and slow enough to avoid evaluating gradients with inadequate expected compound separation.

To encode the compound separation as the model output, we defined a global separation index (GSI). First, a certain number (500 by default) of MS signals with the top ion intensities, termed top signals, were selected. Isotopic ions were excluded. To avoid background ions as top signals, we compared the apex of the chromatographic peak (i.e., peak height) with the average intensity of the peak. We required top signals to have a peak height that is more than double the average intensity by default. The detailed algorithm for selecting top signals is described in **Supplementary Note 10**. With the top signals, we further computed the sum of squared retention time intervals (SQRTI). Suppose that *m* top signals were selected. Their retention times were ranked and concatenated with the boundary of data acquisition time, denoted as *T* = 〈*t*_0_, *t*_1_, …, *t*_*m*+2_〉, where *t*_0_ represents the start of data acquisition (0 min in most cases), and *t*_*m*+2_represents the end of data acquisition. The retention time intervals were then defined as the differences between two adjacent elements in *T*, denoted as *V* = 〈*v*_1_, …, *v*_*m*+1_〉. The SQRTI is given by 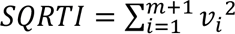. Then, the GSI is given by

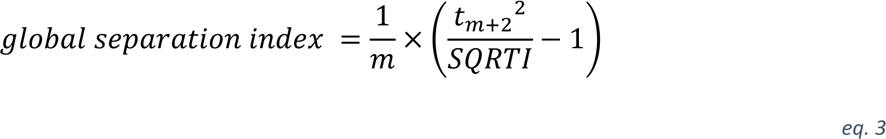

which is a singular value ranging from 0 to 1. The deduction of scaling SQRTI to GSI is detailed in **Supplementary Note 2**.

### Surrogate model

The surrogate model constructs the statistical relationship between the input (i.e., LC gradient) and output data (i.e., GSI) across the entire search space. Gaussian process regression (GPR) was employed to construct the surrogate model for Bayesian optimization, which was implemented in Python using *scikit-learn* package (ver. 1.0.2). The covariance function (i.e., kernel function) in the Gaussian process determines the overall structure of the function distribution, which is a critical hyperparameter in GPR. Here, we utilized Matérn32 kernel to allow for highly flexible experimental data modelling. The covariance function of Matérn32 kernel is given by

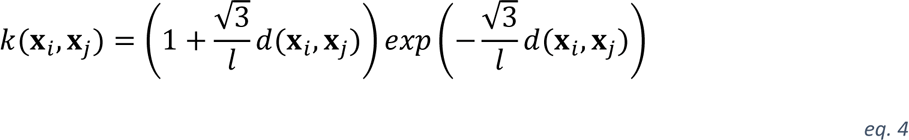

where *d*(*X*_*i*_, *X*_*j*_) is the Euclidean distance between two data points, and *l* is a length-scale parameter (*l* > 0). The Matérn32 kernel allows high flexibility to model the unknown function between the LC gradient and GSI, providing the mean and variance of the posterior distribution in the GPR model. The kernel hyperparameters (e.g., *l*) control the function distribution characteristics, such as smoothness and noise level. When training a GPR model, the kernel hyperparameters are optimized during the fitting process by maximizing the log marginal likelihood, as implemented in *scikit-learn*.

### Acquisition function

The acquisition function determines the LC gradient to be experimentally tested next (**Supplementary Note 4**) for a better experimental outcome. We found that EI demonstrates the best performance as an acquisition function in the gradient optimization problem. With a GPR surrogate model, the improvement function is given by

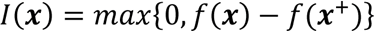

When conditioned in the gradient optimization problem, *f*(X) is the predicted GSI for a given gradient setting X by GPR, and *f*(X^+^) is the best GSI observed in the LC experiment so far. Evaluation of I(X) on a Gaussian posterior distribution yields the expected improvement

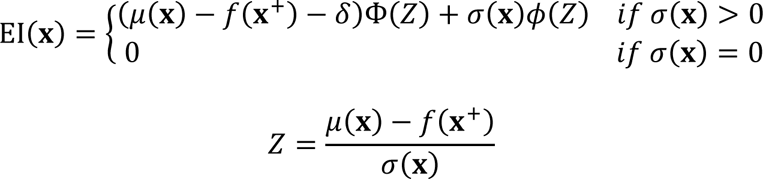

where μ(X) and σ(σ) denote the mean and standard deviation of the posterior distribution at X respectively, and Φ(·) and ϕ(·) denote the cumulative distribution function (CDF) and probability density function (PDF) of the standard normal distribution, respectively. The empirical parameter *δ* was set to 0.01 to balance data exploration and exploitation according to Lizotte’s experiments^24^. The LC gradient to be experimentally tested next is found by searching a gradient setting X in a finite search space that can achieve the largest EI(X). EI was benchmarked with four other acquisition functions, detailed in **Supplementary Note 5**.

### *D. melanogaster* strains and sample collection

The strain used in this study was *w*^1118^ (BDSC 3605), obtained from the Bloomington Stock Center (Bloomington, IN, USA). *Drosophila* stocks were maintained on yeast-sugar-cornmeal food at 25°C in a 12:12 hour light:dark cycle.^47^ Adult *w*^1118^ laid eggs onto grape plates; after 24 hr newly-hatched larvae were transferred to food vials at a density of 50 larvae per 10 mL food (diet consists of 20.5 g sucrose, 70.9 g D-glucose, 48.5 g cornmeal, 45.3 g yeast, 4.55 g agar, 0.5g CaCl2•2H2O, 0.5 g MgSO4•7H2O, and 11.77 mL acid mix (propionic acid/phosphoric acid)). Males and females were separated as late pupae by the presence (males) or absence (females) of sex combs. Pupae were kept in single-sex groups of 20 flies per vial until five days post-eclosion; flies were transferred onto fresh food every two days. Abdomen carcasses were isolated from unmated 5- day-old male and female flies. Each carcass was snap frozen after dissection on dry ice in a 2 mL microcentrifuge tube, and stored at –80°C until metabolome and lipidome extraction. Each biological replicate consisted of abdominal carcasses isolated from 30 flies. A total of 9 biological replicates were collected for each sex.

### Sample preparation and untargeted metabolomics

A total of seven data sets were utilized in this work: human urine metabolomics data with 142 gradient settings, mouse cecum metabolomics data, human urine metabolomics data, human serum metabolomics data, human serum lipidomics data, *Drosophila* abdomen metabolomics data, and *Drosophila* abdomen lipidomics data. Their sample preparation procedures, LC-MS/MS experimental settings, and data processing steps are detailed in **Supplementary Notes 3 and 6-9**. LC-MS analysis was performed on an Impact II ultra-high resolution Qq-time-of-flight mass spectrometer (Bruker Daltonics, Bremen, Germany) coupled with a 1290 Infinity II UHPLC system (Agilent Technologies, Palo Alto, CA, USA). Hydrophilic interaction chromatography (HILIC) separation was performed on a SeQuant ZIC-pHILIC column (150 mm × 2.1 mm, 5 μm, 200 Å) and a SeQuant ZIC-HILIC column (50 mm × 2.1 mm, 5 μm, 200 Å) (MilliporeSigma, Burlington, MA, USA). Reversed phase (RP) separation was achieved on a Waters UPLC Acquity BEH C18 Column (1.0 mm × 100 mm, 1.7 µm, 130 Å, Milford, MA, USA).

### LC-MS data processing

The raw MS data were converted to ABF format in Reifycs Abf Converter (ver. 4.0.0). Then, the converted data were processed in MS-DIAL (ver. 4.90) for chromatographic peak detection, feature alignment, and compound annotation. Only the MS data acquired under the same gradient setting were aligned using MS-DIAL. NIST 20 Tandem Mass Spectral Library (https://www.nist.gov) and the MS/MS database from the MS-DIAL website (ver. 15) were used for compound annotation. The data processing parameters in MS-DIAL were set as follows: MS1 tolerance, 0.01 Da; MS/MS tolerance, 0.05 Da; mass slice width, 0.05 Da; smoothing method, linear weighted moving average; smoothing level, 3 scans; minimum peak width, 5 scans; alignment retention time tolerance, 0.2; alignment *m/z* tolerance, 0.015. The high-quality features were selected according to the previously reported criteria^25^: the average intensity in QC samples is more than twice the intensity of the method blank sample; feature retention time is within the gradient elution time; the relative standard deviation of QC samples intensities is lower than 25%; and the Pearson correlation between MS signal intensities and loading amounts of QC sample is higher than 0.9. The unique MS/MS spectra were selected by grouping MS/MS spectra with a dot product similarity threshold of 0.95. Spectral entropy^27^ values were computed to evaluate the complexity of MS/MS spectra. Alignment of the high-quality metabolic features from original and optimized LC gradient settings was achieved in R.

### Statistical analysis and visualization

Spearman correlation was computed using the R package *stats* (ver. 4.2.0) to explore the relationship between GSI and other properties of metabolic features. Spline fitting was performed in GraphPad Prism 8. The two-dimensional kernel density was calculated in Python using the *seaborn* package (ver. 0.11.2). The two-sided paired Mann-Whitney *U* test was performed in R using the *stats* package (ver. 4.2.1) to obtain *p* values. UMAP was computed using Hiplot (https://hiplot-academic.com/basic/umap). Spectral entropy was computed in R according to the definition by Li’s work^27^. Model fitting results including R^2^ and median absolute error were calculated in Python using the *scikit-learn* package (ver. 1.0.2).

## Data availability

The datasets in this work are summarized in **Supplementary Information**. Raw MS data are available on demand. Source data are provided with this paper (**Supplementary File**).

## Code availability

Code for performing data analysis, Python package, and Windows software is available at https://github.com/HuanLab/bago. Accessibility is declared in **Supplementary Note 11**.

## Acknowledgments

This study was funded by University of British Columbia Start-up Grant (F18-03001), Canada Foundation for Innovation (CFI 38159), and Natural Sciences and Engineering Research Council (NSERC) Discovery Grant (RGPIN-2020-04895). EJR/PB are funded by the Canadian Institutes for Health Research (PJT-153072). Stocks were used from the Bloomington *Drosophila* Stock Center (NIH P40OD018537) and Vienna *Drosophila* Resource Center (VDRC). We also thank Alisa Hui for proofreading this manuscript.

## Author contributions

H.Y. and T.H. designed the study and wrote the manuscript. H.Y. developed the algorithm, Python package, and Windows software. H.Y. performed the LC-MS experiments and data analysis. P. B. and E. J. R. designed the *Drosophila* experiment. P. B. performed animal experiment. All authors discussed the results and contributed to the final manuscript.

## Additional information

**Supplementary information.** The online version contains supplementary material available at

**Correspondence and requests for materials** should be addressed to Tao Huan.

**Peer review information**

**Reprints and permissions information**

---

**Extended Fig. 1.**
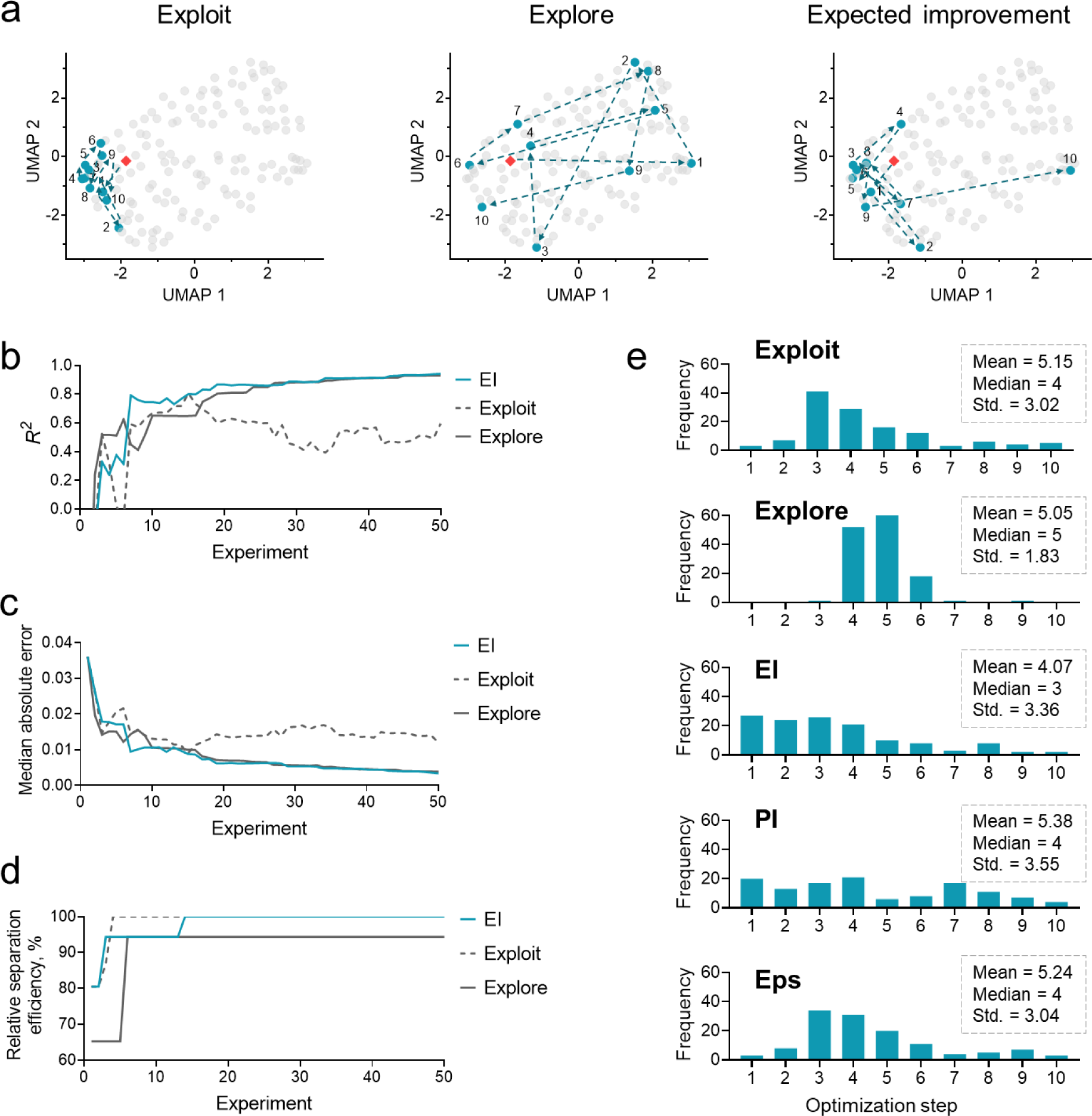
**a**. Three gradient optimization routes differing by acquisition function visualized by uniform manifold approximation and projection (UMAP) plots. The entire search space contains 142 different gradients. Grey dots represent individual LC-MS/MS experiments with unique gradients, and colored dots represent the conducted LC-MS experiment in sequence. Red diamonds represent the initial gradient. **b**, **c**, **d**, Comparison of expected improvement (EI), pure exploration, and pure exploitation on data fitting characterized by R^2^ (**b**), median absolute error (**c**), and improvement of global separation index (**d**). **e**, Histograms to compare the steps required by the five acquisition functions to find an optimal gradient. PI: probability of improvement. Eps: epsilon-greedy.

**Extended Fig. 2.**
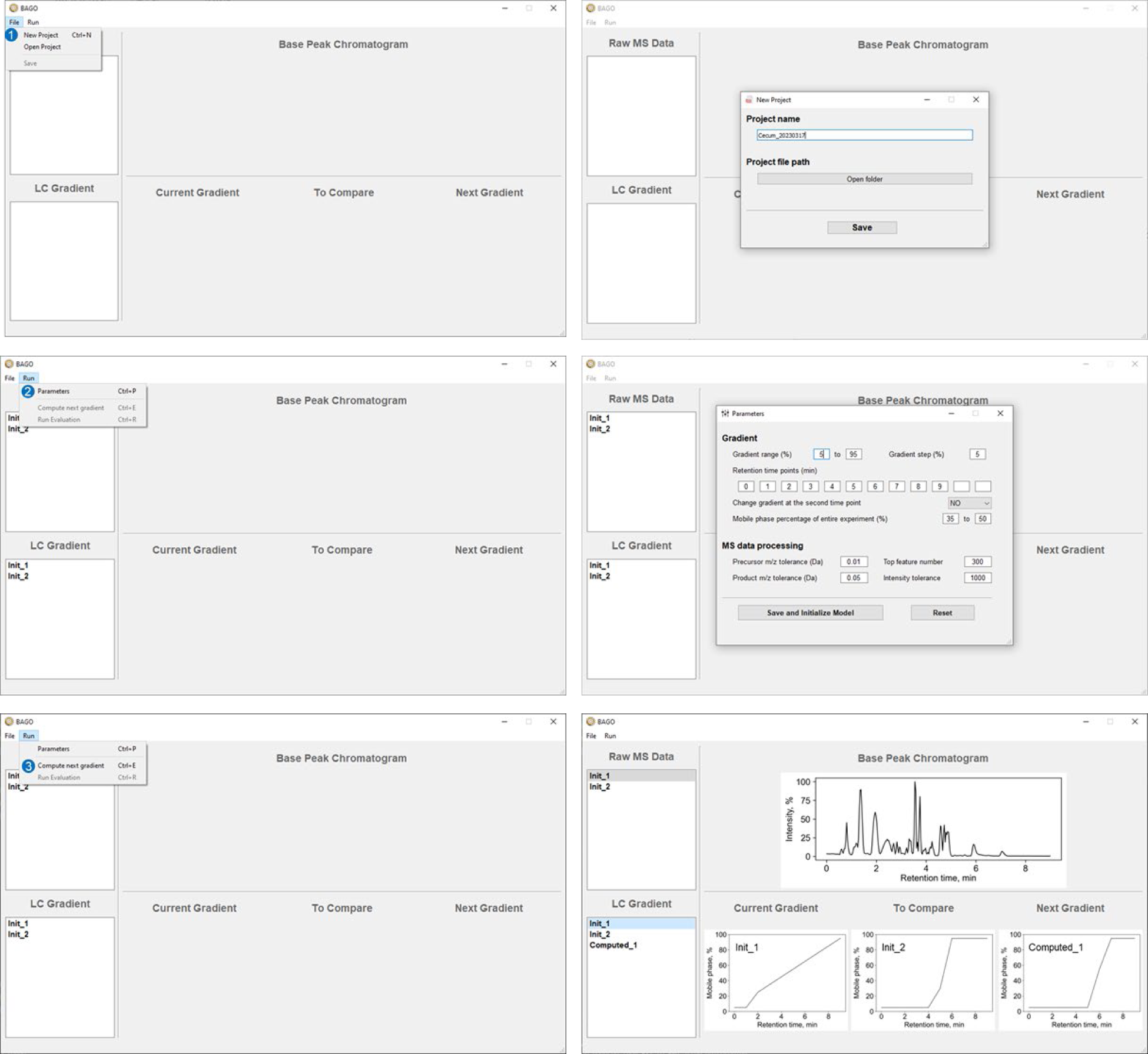
Graphical user interface of BAGO software, including four major panels to manipulate MS data (top left), manipulate LC gradient configurations (bottom left), visualize compound separation by base peak chromatogram (top right), and visualize gradient configurations (bottom right). The entire optimization process has three major steps: create a new project, set parameters, and compute next gradient, as shown on the left column of windows. The windows on the right show the results of each step

