## Supplementary Information for "Bayesian optimization of separation gradients to maximize the performance of untargeted LC-MS"

Internet: https://huan.chem.ubc.ca/

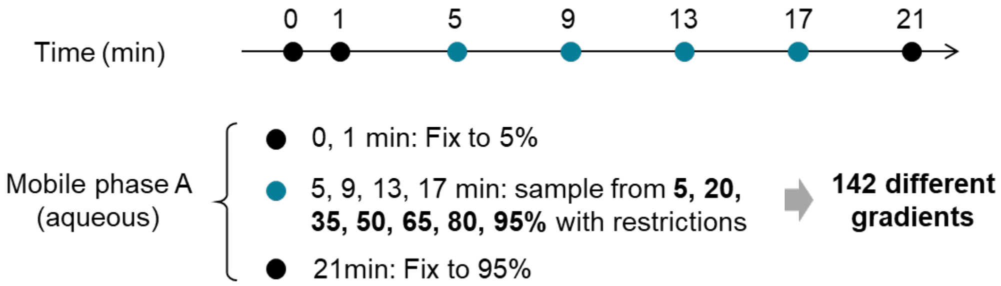

**Supplementary Fig. 1. Illustration of generating LC gradient configurations.** A total of 142 configurations were generated for the human urine metabolomics data set.

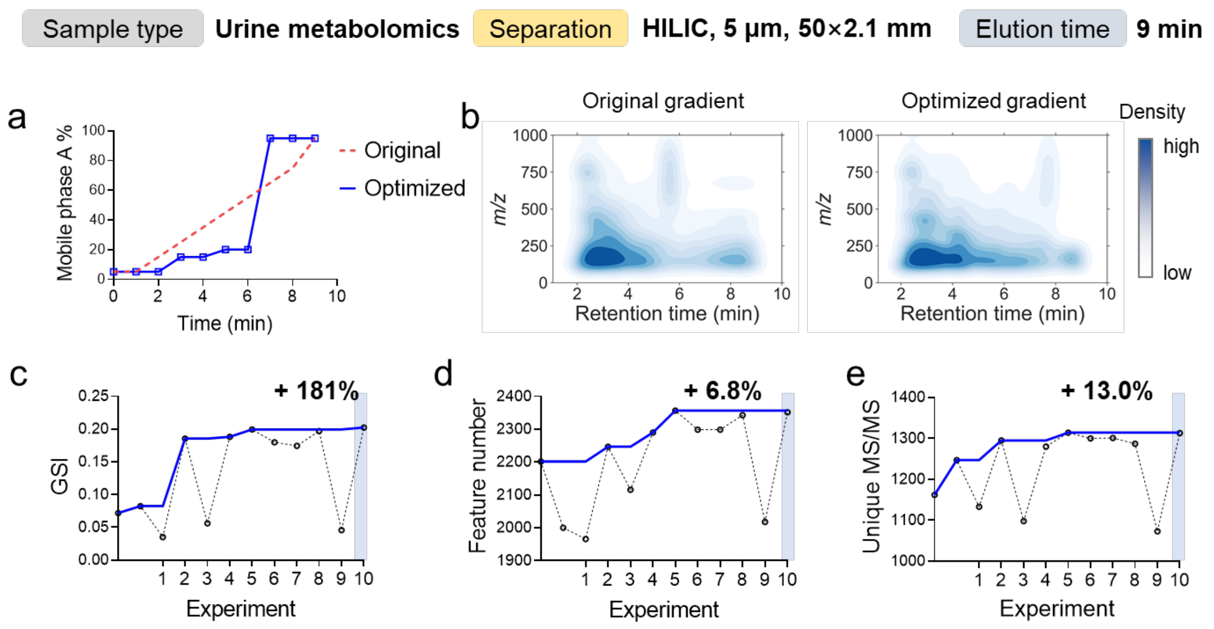

**Supplementary Fig. 2. Gradient optimization of human urine metabolomics sample.** Compound separation was performed on a HILIC LC column with an elution time of 9 minutes. **a**, The optimized LC gradient vs. the original gradient. **b**, Improvement of compound separation visualized by a two-dimensional density plot. **c**, **d**, and **e**, Improved global separation index (GSI) (**c**), total number of metabolic features (**d**), and unique MS/MS spectra during the optimization. The optimal gradient was found on the tenth experiment. Cumulative values are shown as solid curves.

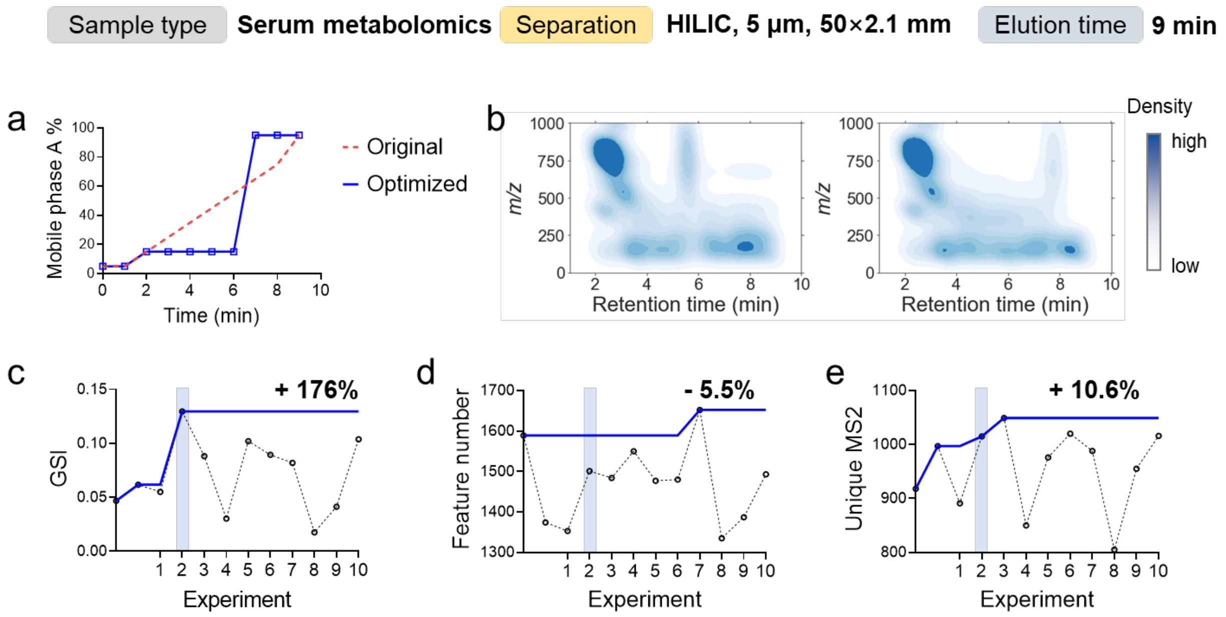

**Supplementary Fig. 3. Gradient optimization of human serum metabolomics sample.** Compound separation was performed on a HILIC LC column with an elution time of 9 minutes. **a**, The optimized LC gradient vs. the original gradient. **b**, Improvement of compound separation visualized by a two-dimensional density plot. **c**, **d**, and **e**, Global separation index (GSI) (**c**), total number of metabolic features (**d**), and unique MS/MS spectra during the optimization. The optimal gradient was found on the tenth experiment. Cumulative values are shown as solid curves.

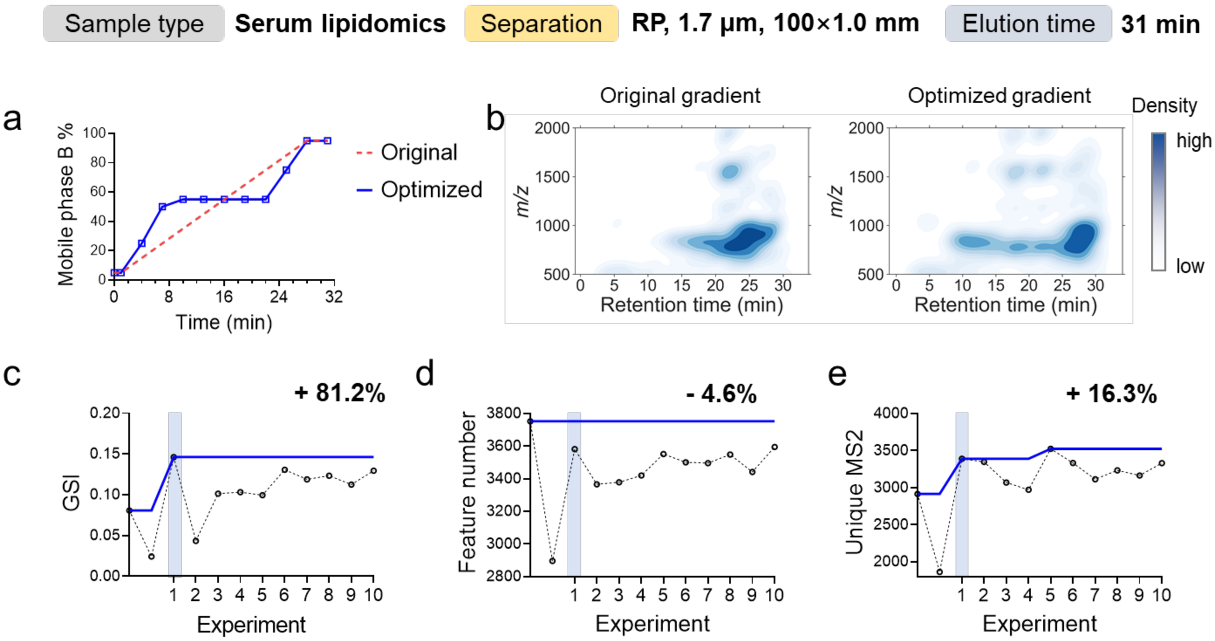

**Supplementary Fig. 4. Gradient optimization of human serum lipidomics sample.** Compound separation was performed on a HILIC LC column with an elution time of 9 minutes. **a**, The optimized LC gradient vs. the original gradient. **b**, Improvement of compound separation visualized by a two-dimensional density plot. **c**, **d**, and **e**, Global separation index (GSI) (**c**), total number of metabolic features (**d**), and unique MS/MS spectra during the optimization. The optimal gradient was found on the tenth experiment. Cumulative values are shown as solid curves.

**Supplementary Note 1. Size of the search space in gradient optimization problems**

Following the principle of gradient design, we define plausible gradient configurations that constitute the entire search space for gradient optimization.

A plausible gradient meets two requirements as described in the **Method** section of the main text. In brief, first, the strong mobile phase percentage should monotonically increase. Second, the total portion of the strong mobile phase should be restricted to a range that ensures the gradient is fast enough for all metabolites to elute and slow enough that users avoid evaluating a gradient with insufficient expected compound separation.

With this definition, we implemented an algorithm in the BAGO software to generate the search space by specifying three parameters:
**Param 1**. the strong mobile phase percentages that can be set (e.g., from 5% to 95% with a 5% interval)
**Param 2**. the time points at which the mobile phase composition can be changed
**Param 3**. the range of % strong mobile phase of total mobile phase used (e.g., from 40% to 60%)

In **Supplementary Fig. 1,** we demonstrated the process of generating a small search space for algorithm development. The principle of this demonstration extends to real gradient optimization problems, except the search space in real application will be much larger.

Here, using the algorithm implemented in the BAGO software (also available as the *bago* python package), we provide two examples with different parameters to generate search spaces.

**Example 1. A 31-min gradient for lipidomics analysis**
**Param 1**. from 5% to 95% with a 5% interval (a total of 19 percentages)
**Param 2**. at 4, 7, 10, 13, 16, 19, 22, 25 and 28 min
**Param 3**. from 45% to 65%
**Search space**: **2,835,542** plausible gradients

**Example 2. A 9-min gradient for metabolomics analysis**

**Param 1**. from 5% to 95% with a 5% interval (a total of 19 percentages)

**Param 2**. at 2, 3, 4, 5, 6, 7 and 8 min

**Param 3**. from 35% to 50%

**Search space**: **261,484** plausible gradients

**Supplementary Note 2. Boundaries of the global separation index**

Global separation index (GSI) is defined to evaluate the global compound separation as a singular value between 0 to 1. It is computed from the sum of squared retention time intervals (SQRTI). This section provides the deduction of the boundaries of GSI and explains why the lower and upper boundaries correspond to the worst and the best separation cases, respectively.

Suppose that $m$ top signals are selected. Their retention times are then ranked and concatenated with the boundary of data acquisition time, denoted as $T = \langle t_0, t_1, \dots, t_{m+2} \rangle$, where $t_0$ represents the start of data acquisition (0 min in most cases), and $t_{m+1}$ represents the end of data acquisition. The retention time intervals are then defined as the differences between two adjacent elements in T, denoted as $V = \langle \Delta t_1, \dots, \Delta t_{m+1} \rangle$. The SQRTI is given by $SQRTI = \sum_{i=1}^{m+1} \Delta {t_i}^2$.

Theoretically, the best separation is achieved when all compounds are equally spaced during the total elution time. Then, we have

$$\Delta t_1 = \Delta t_2 = \cdots = \Delta t_{m+1} = \frac{t_{m+2}}{m+1}$$

*eq. 1*

which gives the lowest SQRTI value according to the Cauchy–Schwarz inequality

$$SQRTI_{min} = \frac{{t_{m+2}}^2}{m+1}$$

*eq. 2*

On the other side, the worst separation is achieved when all compounds elute together; for simplicity, suppose all compounds are eluted out in the beginning. Then, we have

$$SQRTI_{max} = {t_{m+2}}^2$$

*eq. 3*

Combining equations 2 and 3, we have

$$\frac{t^2}{m+1} < \sum_{i=1}^{m+1} \Delta {t_i}^2 < t^2$$

*eq. 4*

From equation 4, we have the equivalent form

$$0 < \frac{1}{m} \times \left( \frac{{t_{m+2}}^{2}}{SQRTI} - 1 \right) < 1$$

eq. 5

with the middle part defined as GSI.

### Supplementary Note 3. Human urine metabolomics data with 142 different gradients

Experimental design

To explore the impact of compound separation on MS-based small molecule profiling, we collected a human urine metabolomics data set consisting of a single urine sample analyzed with 142 different LC gradients (**Supplementary Fig. 1**).

To generate different gradients, we set four time points (5, 9, 13, and 17 minutes) at which the percentage of mobile phase can be changed. Two restrictions were set for plausible gradients, as described in the **Methods** section in the main text. In particular,

1. The percentage of the strong mobile phase is monotonically increasing.
2. The total amount of strong mobile phase used in a gradient ranges from 40-75% to ensure sufficient elution.

We explicitly generated all gradients in a grid search space, yielding 142 different gradients (**Supplementary Fig. 1**). The generated gradients are listed in the **Supplementary File**.

Sample preparation

A human urine sample was collected from a healthy volunteer. The collected biospecimen was stored at -80 °C prior to analysis. For metabolome extraction, 100 µL of urine was mixed with 1000 µL of -20 °C methanol in a 1.5-mL Eppendorf vial and vortexed. The solution was incubated at -20 °C for 2 hours, followed by centrifugation at 14,000 rpm for 15 min at 4 °C. The clear supernatant was dried in a SpeedVac at 20 °C and then reconstituted in 150 µL of acetonitrile and water (1:1, v/v) mixed solvent for LC-MS analysis. A method blank sample was also prepared following the same protocol but without urine.

LC-MS/MS experiment

Hydrophilic interaction chromatography (HILIC) separation was performed on a SeQuant ZIC-pHILIC column (150 mm × 2.1 mm, 5 µm, 200 Å) with an elution time of 21 minutes. Mobile phase A was 5% acetonitrile in water with 10 mM ammonium acetate (Thermo Fisher Scientific) with pH = 4.8, adjusted by acetic acid (MilliporeSigma), and mobile phase B was 5% water in acetonitrile with no buffer. The LC gradients are listed in the **Supplementary File.** The MS was operated in electrospray ionization positive mode (ESI+). The ESI source conditions were set as follows: dry gas temperature, 220 °C; dry gas flow, 7 L/min; nebulizer gas pressure, 1.6 bar; capillary voltage, 4500 V for positive mode. The MS1 analysis was conducted using the following parameters: mass range, 70-1000 *m/z*; spectrum type: centroid, calculated using maximum intensity; absolute intensity threshold: 250. Data-dependent MS/MS analysis parameters: collision energy: 16-30 eV; cycle time, 3 s; spectra rate: 4 Hz when intensity < $10^4$ and 12 Hz when intensity > $10^5$, linearly increased from $10^4$ to $10^5$. External calibration was applied using sodium formate to ensure the *m/z* accuracy before sample analysis. A simple linear gradient was analyzed every 10 injections for quality control purposes.

Data processing

The raw MS data were converted to ABF format in Reifycs Abf Converter (ver. 4.0.0). Then, the converted data were processed in MS-DIAL (ver. 4.90) for metabolic feature extraction[1]. The data processing parameters in MS-DIAL were set as follows: MS1 tolerance, 0.01 Da; MS/MS tolerance, 0.05 Da; mass slice width, 0.05 Da; smoothing method, linear weighted moving average; smoothing level, 3 scans; minimum peak width, 5 scans.

**Supplementary Note 4. Introduction to acquisition functions**

To efficiently find the global maximum, a typical dilemma is to decide whether to explore the regions with high uncertainty (i.e., exploration) or to exploit the regions around the best observation (i.e., exploitation). Exploration maximizes the knowledge that can be gained in the next experiment regarding the objective function, but it may also result in unnecessary effort spent on querying low-yield regions. On the other hand, exploitation typically ensures a promising outcome, but it also risks getting trapped at a local maximum. To balance exploration and exploitation, an acquisition function is created based on GPR. **Fig 1c** shows an acquisition function that comprehensively considers the GPR-predicted objective function and uncertainty, computes the expected improvement over the search space, and provides the most promising LC gradient to be tested in the next LC-MS experiment.

The acquisition function determines the next experimental gradient by balancing data exploration and exploitation. Proper selection of an acquisition function is the key to improve the efficiency of gradient optimization[3]. In this work, we compared five acquisition functions, including pure exploitation, pure exploration, expected improvement, probability of improvement, and epsilon-greedy algorithm.

Pure exploitation

Pure exploitation refers to the selection of the maximum of the estimated objective function within a search space. As an acquisition function, it only focuses on selecting the point with the highest predicted value without considering the uncertainty of the prediction or the potential benefit of exploring other regions of the search space.

Pure exploration

Pure exploration selects the next experiment based on the uncertainty of the prediction without considering the predicted mean of the objective function. It is designed to guide the search on regions where the uncertainty is high, with the goal of obtaining more information about the objective function.

Expected improvement

Expected improvement (EI) measures how much better the candidate point is expected to perform compared to the best observation found by current experiments[4]. As introduced in the main text, with a Gaussian processing (GP) surrogate model, the improvement function is given by

$$\mathrm{I}(\mathbf{x}) = \max\{0, f(\mathbf{x}) - f(\mathbf{x}^+)\}$$

When conditioned in the gradient optimization problem, $f(\mathbf{x})$ is the predicted GSI for a given gradient setting $\mathbf{x}$ by GP regression, and $f(\mathbf{x}^+)$ is the best GSI observed in the LC experiment so far. Evaluation of $\mathrm{I}(\mathbf{x})$ on a Gaussian posterior distribution yields the expected improvement

$$\mathrm{EI}(\mathbf{x}) = \begin{cases} (\mu(\mathbf{x}) - f(\mathbf{x}^+) - \delta)\Phi(Z) + \sigma(\mathbf{x})\phi(Z) & if\ \sigma(\mathbf{x}) > 0 \\ 0 & if\ \sigma(\mathbf{x}) = 0 \end{cases}$$

$$Z = \frac{\mu(\mathbf{x}) - f(\mathbf{x}^+)}{\sigma(\mathbf{x})}$$

where $\mu(\mathbf{x})$ and $\sigma(\mathbf{x})$ denote the mean and standard deviation of posterior distribution at $\mathbf{x}$, respectively, and $\Phi(\cdot)$ and $\phi(\cdot)$ denote the cumulative distribution function (CDF) and probability density function (PDF) of the standard normal distribution, respectively.

Probability of improvement

Probability of improvement (PI) measures the probability that the value of the function being optimized will be improved by a potential candidate point[4]. In particular,

$$\mathrm{PI}(\mathbf{x}) = P(f(\mathbf{x}) \geq f(\mathbf{x}^+))$$

With a GP surrogate model, the PI is given by

$$\mathrm{PI}(\mathbf{x}) = \Phi\left(\frac{\mu(\mathbf{x}) - f(\mathbf{x}^+)}{\sigma(\mathbf{x})}\right)$$

Epsilon-greedy algorithm

The Epsilon-greedy algorithm is a simple strategy to balance exploration and exploitation[4]. In each iteration of optimization, the algorithm randomly decides to explore or exploit the search space based on a predefined probability (i.e., epsilon).

**Supplementary Note 5. Selection of acquisition function**

With the objective to identify the optimal gradient with the shortest evaluation times, we hope to improve the algorithm efficiency. The efficiency of Bayesian optimization is most affected by the acquisition function, which is tasked to balance data exploration and exploitation. Trading off between exploration and exploitation is a critical challenge in the data-driven decision-making process. Conditioned on an LC gradient optimization, this trade-off was demonstrated in **Extended Fig. 1** using the urine metabolomics data set containing 142 different gradients. We first reduced the high-dimensional gradient configurations to a two-dimensional space using uniform manifold approximation and projection (UMAP)[5]. The data points closer together on a UMAP plot are more similar. We tracked the decisions made by different optimization strategies initiated from the same LC gradient in the search space, labeled with a red diamond marker in **Extended Fig. 1**. Pure exploitation began to explicitly search a small region in the lower left after two optimization steps. We anticipated that pure exploitation can easily be trapped at a local maximum and waste experimental efforts. On the other hand, pure exploration traversed the whole search space. It acquired the best knowledge of the unknown function yet risked the low-outcome regions. In principle, a decision can be made by considering the predicted mean and variance by Gaussian process regression (GPR) and draw a candidate gradient most likely to be the best. In doing so, an acquisition function called expected improvement was applied, and we observed that its searching route is between pure exploration and exploitation.

Finding the best gradient in a given search space relies on the global knowledge of the unknown function that we can gain from optimization. Therefore, we compared the data fitting performance of the three abovementioned acquisition functions after certain optimizations. Overall, expected improvement and exploration have similar and high coefficients of determination ($R^2$). Both acquisition functions achieve $R^2 > 0.8$ by mid-experiment. In comparison, pure exploitation showed much lower $R^2$, increasing to 0.72 at the 25$^{th}$ step and decreasing to 0.42-0.44 near the end. Pure exploitation also showed significantly higher median absolute error than the other two, indicating poor global knowledge and agrees with the $R^2$ results.

Besides the expected improvement, other acquisition functions, including epsilon-greedy and probability of improvement, were also routinely utilized to balance exploration and exploitation. To assess the performance of different acquisition functions, we compared the number of steps they required to obtain an optimal gradient, defined as gradients with the top three separation efficiencies. Our results showed that, when starting from the same linear gradient and another gradient in the search space, expected improvement requires the fewest median steps to find an optimal gradient. Therefore, the remarkable performance of expected improvement demonstrates its suitability as the default acquisition function in Bayesian optimization of LC gradients.

**Supplementary Note 6. Human urine metabolomics data set**

Experimental design

A human urine sample was utilized to demonstrate the performance of the Bayesian gradient optimization workflow. Ten optimizations were carried out after the original gradient. Global separation index, number of metabolic features, and number of unique MS/MS spectra were examined to evaluate the optimization performance.

Sample preparation

Refer to “Sample preparation” in **Supplementary Note 3**.

LC-MS/MS experiment

Hydrophilic interaction chromatography (HILIC) separation was performed on a SeQuant ZIC-HILIC column (50 mm × 2.1 mm, 5 µm, 200 Å) with an elution time of 9 minutes. Mobile phase A was 5% acetonitrile in water with 10 mM ammonium acetate (Thermo Fisher Scientific) with pH = 9.8, adjusted by ammonium hydroxide (MilliporeSigma), and mobile phase B was 5% water in acetonitrile with no buffer. The LC gradients are listed in the **Supplementary File**. The MS configurations were set as described in section “LC-MS/MS experiment” in **Supplementary Note 3**.

Data processing

As described in **Supplementary Note 3**.

Results

Results are shown in **Supplementary Fig 2**. We began the optimization by generating a large search space that contains 261,484 plausible gradients. A ten-round optimization was carried out after two initial gradients. The best gradient was the tenth experiment, improving the GSI from 0.0718 to 0.202. With the optimized gradient, we profiled 6.8% more metabolic features (from 1660 to 1894) and detected 13.0% more unique MS/MS spectra (from 1162 to 1313).

**Supplementary Note 7. Human serum metabolomics data set**

Experimental design

A human serum sample was utilized to demonstrate the performance of the Bayesian gradient optimization workflow. Ten optimizations were carried out after the original gradient. Global separation index, number of metabolic features, and number of unique MS/MS spectra were examined to evaluate the optimization performance.

Sample preparation

The human serum sample was purchased from Sigma-Aldrich (St. Louis, MO, USA). The sample was stored at -80 °C prior to analysis. For metabolome extraction, 50 µL of serum was mixed with 1000 µL of -20 °C methanol in a 1.5-mL Eppendorf vial and vortexed. The solution was incubated at -20 °C for 6 hours, followed by centrifugation at 14,000 rpm for 15 min at 4 °C. The clear supernatant was dried in a SpeedVac at 20 °C and then reconstituted in 150 µL of acetonitrile and water (1:1, v/v) mixed solvent for LC-MS analysis. A method blank sample was also prepared following the same protocol but without serum.

LC-MS/MS experiment

Hydrophilic interaction chromatography (HILIC) separation was performed on a SeQuant ZIC-HILIC column (50 mm × 2.1 mm, 5 µm, 200 Å) with an elution time of 9 minutes. Mobile phase A was 5% acetonitrile in water with 10 mM ammonium acetate (Thermo Fisher Scientific) with pH = 9.8, adjusted by ammonium hydroxide (MilliporeSigma), and mobile phase B was 5% water in acetonitrile with no buffer. The LC gradients are listed in the **Supplementary File**. The MS configurations were set as described in the section "LC-MS/MS experiment" in **Supplementary Note 3**.

Data processing

As described in **Supplementary Note 3**.

Results

Results are shown in **Supplementary Fig 3**. We began the optimization by generating a large search space that contains 261,484 plausible gradients. A ten-round optimization was carried out after two initial gradients. The best gradient was the tenth experiment, improving the GSI from 0.0468 to 0.130. With the optimized gradient, we detected 10.6% more unique MS/MS spectra (from 1162 to 1313). Notably, 5.5% fewer metabolic features (from 1589 to 1501) were extracted by MS-DIAL in this data set.

**Supplementary Note 8. Human serum lipidomics data set**

Experimental design

A human serum sample was utilized to demonstrate the performance of the Bayesian gradient optimization workflow. Ten optimizations were carried out after the original gradient. Global separation index, number of metabolic features, and number of unique MS/MS spectra were examined to evaluate the optimization performance.

Sample preparation

The same human serum sample was used as described in **Supplementary Note 7**. For lipidome extraction, 50 µL of serum was mixed with 270 µL of -20 °C methanol in a 1.5-mL Eppendorf vial and vortexed. The solution was incubated at -20 °C for 6 hours, followed by adding 900 µL methyl tert-butyl ether (MTBE) for lipid extraction. The one-phase solvent was shaken for 2 min, and 265 µL of water was added to induce the phase separation. After mixing, the solution rested at room temperature for 10 min. Next, the solution was centrifuged at 14,000 rpm for 15 min at 4 °C. The clear upper layer was transferred to a new vial and dried in a SpeedVac at 20 °C. The sample was then reconstituted in 150 µL of acetonitrile and isopropanol (1:1, v/v) mixed solvent for LC-MS analysis. A method blank sample was also prepared following the same protocol but without serum.

LC-MS/MS experiment

Reversed phase (RP) separation was achieved on a Waters UPLC Acquity BEH C18 Column (1.0 mm ×100 mm, 1.7 µm, 130 Å, Milford, MA, USA) with an elution time of 31 minutes. Mobile phase A was acetonitrile and water (6:4, v/v) with 2 mM ammonium formate (pH = 4.8, adjusted by formic acid), and mobile phase B was isopropanol and acetonitrile (9:1, v/v). The LC gradients are listed in the **Supplementary File.** The MS was operated in electrospray ionization positive mode (ESI+). The ESI source conditions were set as follows: dry gas temperature, 220 °C; dry gas flow, 7 L/min; nebulizer gas pressure, 1.6 bar; capillary voltage, 4500 V for positive mode. The MS1 analysis was conducted using following parameters: mass range, 70-1000 *m/z*; spectrum type: centroid, calculated using maximum intensity; absolute intensity threshold: 250. Data-dependent MS/MS analysis parameters: collision energy: 16-30 eV; cycle time, 3 s; spectra rate: 4 Hz when intensity < $10^4$ and 12 Hz when intensity > $10^5$, linearly increased from $10^4$ to $10^5$. External calibration was applied using sodium formate to ensure the *m/z* accuracy before sample analysis.

Data processing

As described in **Supplementary Note 3**.

Results

Results are shown in **Supplementary Fig 4**. We began the optimization by generating a large search space that contains 2,835,542 plausible gradients. A ten-round optimization was carried out after two initial gradients. The best gradient was the tenth experiment, improving the GSI from 0.0806 to 0.146. With the optimized gradient, we detected 16.3% more unique MS/MS spectra (from 2916 to 3390). Notably, 4.6% fewer metabolic features (from 3753 to 3582) were extracted by MS-DIAL in this data set.

**Supplementary Note 9. Biological application**

Experimental design

Parallel metabolomics and lipidomics analyses of *Drosophila* abdomen were performed to study the different responses of males and females on a high-sugar diet. Ten optimizations were carried out after the original metabolomics gradient and the original lipidomics gradient. Global separation index, number of metabolic features, number of unique MS/MS spectra, annotated metabolites, and annotated lipids were examined to evaluate the optimization performance.

Sample preparation

A total of 180 *Drosophila melanogaster* (90 females and 90 males) abdomen regions were harvested for this biological application. For each sex, the 90 individuals were divided into 9 groups with 10 individuals per sample. Samples were subjected to a dual-phased extraction. First, *Drosophila* abdomen samples were homogenized with a Mini-Beadbeater-24 (Biospec Products, Bartlesville, United States) at 3800 rpm. A total of 333 μL 90% MeOH was added to the *Drosophila* abdomen sample in a 2.0 mL screw cap microcentrifuge tube, which was purchased from Thermo Fisher Scientific (Waltham, MA). Two 15-second homogenization cycles were performed with a 1-min rest on dry ice in between. The homogenized solution was mixed with 1000 μL of MTBE and shaken for 5 min. Then, 317 μL of water was added to induce the phase separation. After mixing, the solution rested at room temperature for 10 min. Next, the two-layer solution was centrifuged at 14,000 rpm for 15 min at 4 °C. The clear upper and lower layers were separated to new vials, dried in a SpeedVac at 20 °C, and reconstituted in 70 μL of acetonitrile and isopropanol (1:1, v/v) and 70 μL of acetonitrile and water (1:1, v/v) mixed solvent for LC-MS analysis, respectively. A method blank sample was also prepared following the same protocol but without *Drosophila* abdomen sample.

LC-MS/MS experiment

Instrumental configuration was set as per **Supplementary Note 7** for metabolomics and as **Supplementary Note 8** for lipidomics. The LC gradients are listed in the **Supplementary File**.

Data processing

The raw MS data were converted to ABF format in Reifycs Abf Converter (ver. 4.0.0). The converted data were processed in MS-DIAL (ver. 4.90) for feature extraction. The data processing parameters in MS-DIAL were set as follows: MS1 tolerance, 0.01 Da; MS/MS tolerance, 0.05 Da; mass slice width, 0.05 Da; smoothing method, linear weighted moving average; smoothing level, 3 scans; minimum peak width, 5 scans.

The generated feature intensity tables for metabolomics and lipidomics were used for quantitative comparison. First, the *MAFFIN* R package[2] was used to select high-quality metabolic features, correct MS signal intensities, and normalize total sample amounts. Then, compound annotation was achieved using the internal library in MS-DIAL for lipidomics. For metabolomics, compound annotation was performed using the NIST 20 library.

**Supplementary Note 10. Top signal selection**

To compute global separation index (GSI), a certain number of MS signals with the highest intensities (i.e., top signals) are selected. We select MS signals from unique compounds with good chromatographic peak shapes by establishing multiple orthogonal criteria. In general, there are three steps to the selection of top signals, including preselection, purification, and deisotoping.

<u>Preselection of top signals</u>

**Require**: MSData (MS data ordered as scan number), s (number of top signals to be selected)

1: Record all MS signals (i.e., *m/z* and intensity) in the first MS1 scan to a top signal list to initiate the selection process.

2: **for** *i* in 2, 3, …, N (total scan number) **do**

3: check **if** a MS signal in current scan also shows in last scan

4: if **yes**, compare their intensities and keep the larger one

5: if **no**, add it to the top signal list

6: end **for**

7: rank the top signals by intensity and select 2×s top signals for the next step

<u>Purification of top signals</u>

**Require**: topSignals (a list of pre-selected top signals)

1: **for** *i* in 1, 2, …, 2×s (s is the required top signals) **do**

2: check **if** the peak height is larger than twice the average intensity of the feature

3: if **yes**, pass

4: if **no**, remove from the list

5: end **for**

<u>Deisotoping</u>

**Require**: topSignals (a list of purified top signals)

1: **for** *i* in 1, 2, …, l (l is the number of purified top signals) **do**

2: check **if** any other top signal can be an isotope of the current ion by matching retention time and *m/z*

```
        if **yes**, remove the isotope with lower intensity
end **for**
```

**Supplementary Note 11. Accessibility**

The proposed Bayesian optimization workflow and framework for gradient optimization can be accessed from a Python application programming interface (API) and a Windows application. The proposed tools enable highly efficient gradient optimization, omics-scale evaluation on compound separation, and broader discovery of chemical space.

Python package, *bago*

*bago* is a Python package for Bayesian optimization of liquid chromatographic elution gradients. *bago* enables flexibility, customization, and extension to the proposed workflow. More information on *bago* can be found at

**Download**: https://pypi.org/project/bago
**Documentation**: https://bago.readthedocs.io/en/latest
**Source code**: https://github.com/Waddlessss/bago
**Bug reports**: https://github.com/Waddlessss/bago/issues

Windows software, BAGO

BAGO is a Windows Application in .exe format for Bayesian optimization of liquid chromatographic elution gradients. BAGO has a clear and simple user interface and requires no coding experience. More information on the BAGO Windows Application can be found at

**Download and Documentation:**
**Address 1:** https://github.com/Waddlessss/bago/releases
**Address 2:** https://1drv.ms/f/s!AoPvCNx85weikRIJQyzN8RHOPvRO?e=lgsl3p
**Bug reports**: https://github.com/Waddlessss/bago/issues
